# Consistent, scale-dependent differences in the biogeography of host-associated and free-living microbiomes across systems

**DOI:** 10.64898/2026.04.16.718993

**Authors:** Joshua H. Dominguez, Andreas Härer, Christopher B. Wall, Diana J. Rennison, Celia C. Symons, Jonathan B. Shurin

**Affiliations:** School of Biological Sciences, Department of Ecology, Behavior, & Evolution, University of California San Diego, La Jolla, California, USA; Department of Earth Sciences, University of Hawai‘i at Mānoa, Honolulu, Hawai‘i, USA; Department of Ecology and Evolutionary Biology, University of California, Irvine, California, USA

**Author notes:** Corresponding author: Joshua H. Dominguez.

**Keywords:** bacterioplankton, salmonids, stickleback, lakes, biogeography, metacommunities

## Abstract

Microbial communities are critical to the functioning of ecosystems and shape the ecology and evolution of host organisms. However, we have a limited understanding of how host-associated and free-living microbes differ in their structure and biogeography. Here, we test whether host-associated (fish gut) and free-living (lake bacterioplankton) microbes exhibit different metacommunity structure, spatial turnover, and consistency with neutral expectations using two independent lake systems. We characterized microbial communities in lake water (Vancouver Island and Sierra Nevada) and guts in two fish species (stickleback and brook trout) using 16S amplicon sequencing. We compared alpha and beta diversity within lakes, quantified spatial turnover (distance-decay), and tested for departure from neutral abundance-occurrence expectations between bacterioplankton and fish gut microbiomes. Fish microbiomes had lower alpha diversity compared to bacterioplankton, but higher beta diversity within lakes. Bacterioplankton were more similar across lakes yet showed stronger patterns of spatial turnover with distance than fish gut microbiomes. A neutral model explained a substantial proportion of abundance-occurrence relationships in bacterioplankton communities but performed poorly for fish-associated microbes. Our study indicates that host-associated and free-living microbes have disparate patterns of metacommunity structure and spatial turnover consistent with differences in the strength of neutral ecological processes. Fish microbiomes were less diverse at the local scale but more variable across space and time than bacterioplankton communities, suggestive of potentially strong local selection and/or reduced microbial exchange among hosts compared to environmental communities. Importantly, we observed highly consistent patterns across both lake systems despite differences in host species, sampling design, and region, demonstrating that differences in the distribution of host and environmental microbes are potentially widespread.

This study demonstrates how host association fundamentally alters the diversity and spatial distribution of microbes, emphasizing the need to incorporate hosts into broader frameworks of microbial biogeography.

## 1. INTRODUCTION

Microorganisms display many of the same patterns of diversity as plants and animals including species-area relationships (Horner-Devine et al. 2004), distance-decay patterns (Martiny et al. 2011), and elevation diversity relationships (Nottingham et al. 2018). These patterns emerge from the same fundamental processes of speciation, immigration, environmental selection, extinction, and drift (Xu et al. 2020; Dickey et al. 2021; Härer & Rennison 2023). Yet the strength of these processes can vary between microorganisms and macroorganisms, leading to differences in the magnitude of observed patterns (Horner-Devine et al. 2004). While these differences are increasingly documented, less is known about how patterns vary among microbial communities occupying distinct habitats. Microbes occur as free-living organisms in the environment and as symbionts associated with eukaryotic hosts, which impose distinct dispersal pathways and selective pressures. Consequently, host-associated and free-living microbial communities may exhibit contrasting diversity and metacommunity structure across environments.

Most research has focused on free-living microbes in aquatic or soil environments; however, a growing body of work documents the distribution and diversity of microbes associated with plant and animal hosts (Neu et al. 2021; Neu et al. 2021; Henry & Ayroles 2022; Härer & Rennison 2023; Mannochio-Russo et al. 2023). Microbial symbionts form close associations with host organisms and can significantly influence host ecology and evolution by altering immune function (Hooper et al. 2012), metabolism (Utzschneider et al. 2009), and life history (Macke et al. 2017). Similar to free-living microbes, host-associated microbiomes are shaped by a combination of stochastic and deterministic factors including dispersal limitation (Moeller et al. 2017), priority effects (Debray et al. 2022), and competition (Coyte et al. 2015). Despite horizontal transmission between host-associated and free-living assemblages, these communities are often taxonomically and functionally divergent (Cuellar-Gempeler & Leibold 2019; Härer et al. 2020). Consequently, it is critical to understand how diversity patterns, biogeography, and potential assembly processes vary between host-associated and free-living microbial communities as both shape ecosystem processes at different scales.

Variation in dispersal among habitat patches may contribute to differences in diversity patterns between host-associated and free-living microbes. Theory suggests that dispersal rate among habitat patches influences local (alpha), regional (gamma), and between-community (beta) components of diversity (Holt 1993; Loreau & Mouquet 1999). Increasing dispersal rates can enhance local diversity via colonization from neighboring habitat patches but reduce regional or beta diversity as communities become more homogenized (Mouquet & Loreau 2003). Early research assumed that microbial dispersal was global and that environmental selection governed their assembly across environmental gradients exclusively (Finlay et al. 2002; Fenchel & Finlay 2004; Wit et al. 2004; Finlay & Esteban 2007). However, microbes exhibit patterns of diversity and distribution that suggest dispersal rates vary across space, time, and environments (Echenique-Subiabre et al. 2025). Microbes closely associated with hosts may experience stronger dispersal limitation than free-living microbes, as host association may restrict opportunities for transmission and exchange among habitat patches. Consequently, slower dispersal may reduce local diversity and drive higher beta diversity amongst individual hosts. In contrast, free-living microbes may disperse more freely across landscapes and environments through aerosols, dust, or hydrological connectivity resulting in higher alpha and lower beta diversity.

Ecological selection may also differ between host-associated and free-living microbes. Hosts exert strong selection on the structure of microbiomes mediated through an array of factors including physiology (Fontaine & Kohl 2020; Procházková et al. 2024), diet (Kartzinel et al. 2019; Youngblut et al. 2019), and genetic architecture (Spor et al. 2011; Smith et al. 2015; Frankel-Bricker et al. 2020). In particular, gut microbiome diversity is shaped by phylogenetic and gut morphological differences among hosts (Reese & Dunn 2018). Variation in these host-scale selection mechanisms may drive higher species-turnover in microbiomes, especially among individuals in different environments or with divergent evolutionary histories. In contrast, selection in free-living microbes is driven by environmental gradients and temporal seasonality that can alter diversity patterns (Gilbert et al. 2012; Signori et al. 2014; She et al. 2021). Despite being connected in the same systems, divergence in the scale and strength at which ecological selection acts upon host and free-living microbes may differentially shape microbial diversity patterns and metacommunity structure.

Differences in dispersal and selection may also drive varying spatial turnover between host-associated and free-living microbiomes. Spatial turnover (distance-decay of similarity) is a pattern where communities become less similar with increasing distance due to neutral dispersal limitation or species-sorting (Tilman 1982; Hubbell 2001; Chave 2004; Alonso et al. 2006; Soininen et al. 2007; Wirens 2011). In microbial communities, patterns of spatial turnover are increasingly documented, but their magnitude varies considerably across study systems (Clark et al. 2021), habitats (Delleuze et al. 2024), and spatial scales (Martiny et al. 2011). Recent work suggests that host-associated microbes show reduced species-turnover compared to free-living microbes across similar spatial scales (Clark et al. 2021; Suzzi et al. 2023). This pattern may reflect stronger dispersal limitation and/or ecological selection among host-associated microbes, particularly for those residing within the digestive tracts of animals. Nevertheless, the extent to which host-associated and free-living microbes differ in spatial turnover across hosts and environments remains underexplored.

Metacommunity theory and neutral models provide useful frameworks for conceptualizing microbial community dynamics across hosts, environments, and space (Leibold et al. 2004; Economo & Keitt 2008; Miller et al. 2018). Neutral models predict that the occurrence of taxa across habitat patches is driven primarily by stochastic dispersal and drift, resulting in predictable relationships between abundance and occurrence (Sloan et al. 2006; Woodcock et al. 2007). Deviation from neutral expectations suggest the influence of ecological selection or non-neutral dispersal ability. In addition, neutral models can identify microbial taxa under putative selection and those that make up the “core” microbiome of a given environment or host (Hernandez et al. 2017; Neu et al. 2021). Therefore, comparing neutral model fits and diversity patterns between host-associated and free-living microbial communities can provide insights into the relative importance of dispersal and selection across habitats.

This study asks whether free-living and host-associated microbes exhibit differences in alpha or beta diversity, spatial turnover (distance decay), or fit to neutral model expectations. We compared the metacommunity structure of fish gut and lake water (bacterioplankton) microbiomes across two independent lake systems over multiple years. In mountain lakes of the Sierra Nevada, CA, USA, we sampled fecal microbiomes of brook trout (*Salvelinus fontinalis* (Mitchill, 1814)) and free-living water microbes from fifteen lakes that were repeatedly sampled across multiple years. On Vancouver Island, BC, Canada, we collected threespine stickleback (*Gasterosteus aculeatus* (Linnaeus, 1758)) gut and water microbiomes from nineteen post-glacial lakes that were sampled at one time point across three years. We tested whether fish gut microbiomes and bacterioplankton exhibit different patterns of spatial turnover, diversity within and among lakes, and are differentially shaped by neutral processes using an abundance-occurrence neutral model. By leveraging these two systems, we aim to uncover whether host and free-living microbiomes display metacommunity patterns that suggest differences in the roles of deterministic and neutral community assembly processes.

## 2. MATERIALS AND METHODS

### 2.1 Lake and fish sampling

Detailed sampling methods are described in Supporting Information Methods 1. Briefly, we sampled brook trout fecal and free-living water (bacterioplankton) microbiomes from fifteen Sierra Nevada (abbreviated SN) lakes spanning an 800m elevation gradient across multiple years (Table S1). Geographic distances between lakes ranged from 0.57 to 35 km. Bacterioplankton from the same fifteen lakes were repeatedly sampled once annually across the summer seasons (late June – early September) of 2016, 2017, 2019, 2021, and 2022 (63 total samples; Table S1) with 0.22 μm Sterivex filter cartridges (Millipore) using previously established protocols (SI Methods 1; Schulhof et al. 2020; Wall & Perreault et al. 2025). In parallel, we collected fecal material from 186 brook trout (n = 5 – 20 individuals per lake [n = 14 – 15 lakes per year]) from the same lakes during the summer seasons of 2021 and 2022 (Table S1). All sampling was performed under National Park Service permit YOSE-2022-SCI-0031, California Department of Fish and Wildlife (CDFW) permit GF-211970003-21197-001, and UCSD IACUC Protocol #S14140.

On Vancouver Island (abbreviated VI), we sampled bacterioplankton and stickleback-gut microbiomes from nineteen lakes at one time point per lake across a geographic range from 1 to 81 km during the spring/early summers of 2020, 2021, and 2022 (Table S1). We sampled bacterioplankton by hand-filtering lake water collected from the shore using a bleach sterilized syringe equipped with a 0.2 µm filter (Whatman plc, Maidstone, UK). At each lake we collected 3 – 4 replicate filters totaling 74 water samples. Simultaneously, we collected whole intestines from 541 stickleback (n = 18 – 30 individuals per lake) that were captured using unbaited minnow traps under British Columbia collection permits NA20-602264, MRVI21-619908, and NA22-713085.

### 2.3 DNA extraction, PCR, and sequencing

See supporting information for detailed documentation on sample and library preparation (Supporting Information Methods 2). DNA was extracted from brook trout, stickleback, and VI lake water samples using the Qiagen DNeasy PowerSoil Pro Kit (Qiagen, cat no. 47014). For SN sterivex filters we used the Qiagen DNeasy PowerWater Kit (Qiagen, cat no. 14900-100-NF). To characterize microbial communities, we amplified the V4 region of the 16S ribosomal RNA gene using barcoded 515F and 806R primer sets (https://github.com/SchlossLab/MiSeq_WetLab_SOP/; Apprill et al. 2016; Parada et al. 2016). Amplified DNA was sequenced on the Illumina MiSeq platform (PE250 for SN samples, PE300 for VI).

### 2.4 Bioinformatic processing

We trimmed primers and adapters from raw reads (fastq files) using ‘cutadapt’ v4.3 (Martin 2011) and processed reads through the DADA2 pipeline (Callahan et al. 2016) in R version 4.1.1 (R Core Team 2025). Forward and reverse reads were quality inspected and subsequently truncated at 225 and 200 base pairs respectively. Truncated reads were merged and resolved to amplicon sequence variants (ASVs). Chimeric sequences were removed, and we assigned taxonomy to ASVs using the 2021 SILVA 138.1 16S rRNA sequence database (Quast et al. 2013). Extraction controls were used to remove possible contaminants using the ‘decontam’ package (Davis et al. 2018). With the ‘phyloseq’ package (McMurdie & Holmes 2013), we removed samples with less than 100 reads and filtered out ASVs that were assigned to chloroplast, mitochondria, or were in less than two samples. Prior to downstream analyses we normalized ASV tables to 3000 reads using scaling with ranked subsampling with the ‘SRS’ R package (Beule & Karlovsky 2020; Heidrich et al. 2021). After normalizing and removing samples with less than 3000 reads, our final dataset consisted of 788 samples (469 stickleback, 169 brook trout, 61 SN lakes, 71 VI lakes) with 21,516 ASVs. All downstream analyses were performed using R Version 4.4.2 (R Core Team, 2025).

### 2.5 Microbiome composition

To account for regional and spatiotemporal differences in sampling design, we analyzed samples from the SN and VI separately. We calculated Bray-Curtis distances on Hellinger-transformed ASV count matrices using *vegdist()* (‘vegan’ package; Oksanen et al. 2007), and phylogenetic differences were estimated using weighted Unifrac distances (*distance()* function in ‘phyloseq’).

We used permutational multivariate analysis of variance (PERMANOVA; *adonis2()* in ‘vegan’) to examine the effects of habitat type (fish gut vs bacterioplankton), lake, and year. In each system, we fit two complementary PERMANOVA models to independently test for spatial and temporal changes in community structure. In the spatial model, habitat type, lake, and their interaction were included as fixed effects, and permutations were constrained within years (strata argument in *adonis2()*) to test for spatial variation while controlling for time. In the temporal model, habitat type, sampling year, and their interaction were fixed effects, and permutations were restricted by lake to account for space. This approach allowed us to evaluate how space and time independently shape microbial communities while accounting for repeated sampling.

To examine whether unequal sample size between fish and bacterioplankton influenced our PERMANOVA results, we performed a balanced subsampling sensitivity analysis. Within each lake and year, we randomly subsampled fish to match bacterioplankton sample sizes and repeatedly refit PERMANOVA models using the same structure. We repeated this procedure 100 times and recorded the median effect sizes (R^2^) and significance of terms across iterations to examine the robustness of our results (Table S8).

We identified taxa (Family level) enriched in fish and bacterioplankton/water within each system using Analysis of Compositions of Microbiomes with Bias Correction (ANCOM-BC) on ASV counts with the ANCOMBC package (Lin & Peddada 2020). ANCOM-BC was performed for each system independently to test whether taxonomic contrasts between fish hosts and the environment were similar across study systems.

### 2.6 Differences in Alpha and Beta diversity

Within each study system, we tested whether fish gut and lake water microbiomes exhibited differences in alpha diversity (richness and Shannon) using generalized linear mixed effects models (GLMMs) with the ‘glmmTMB’ R package. Alpha diversity metrics were calculated using *estimate_richness()* in ‘phyloseq’ (McMurdie & Holmes 2013). In both systems, we modeled Shannon diversity and ASV richness as a function of habitat type (fish gut or water). ASV richness was modeled using a negative binomial distribution with a log-link to accommodate over-dispersed count data, and Shannon diversity was modeled with a Gaussian distribution. To account for differences in sampling design, we used different random effects structures for each system. Models for VI samples included random intercepts for year and lake to account for multiple samples collected within lakes during single years. For the SN, where lakes were repeatedly sampled across multiple years, we included random intercepts for lake and year nested within lake (lake/year) to account for repeated sampling through time. This approach allowed us to compare alpha diversity between host-associated and free-living communities while accounting for a hierarchical sampling structure and uneven sample sizes.

We tested for differences in beta diversity (species turnover) between fish gut and bacterioplankton using Bray-Curtis dissimilarities calculated on Hellinger-transformed ASV counts. Due to differences in sampling design, we used distinct measures of turnover for each study system.

For VI samples, we asked whether habitat types differed in within-lake spatial beta diversity (i.e., within-lake sample heterogeneity). We quantified within-lake spatial beta diversity by calculating Bray-Curtis dissimilarities between each sample and the lake-level centroid community (i.e., mean) for each habitat type (fish gut and bacterioplankton) collected on the same date. We defined lake-level centroid communities as the mean ASV abundances across replicate samples within each lake for both habitat types. Distances from centroid communities were averaged across samples to generate lake-level estimates of spatial turnover for each habitat types. Using these lake-level means, we fit a GLMM with a beta distribution (logit-link) predicting within-lake spatial beta diversity as a function of habitat type, with lake and year as random effects.

For SN samples, we asked whether brook trout gut microbes and bacterioplankton differed in within-lake temporal beta diversity by calculating Bray-Curtis distances between samples collected in 2021 and 2022 within each lake. Since brook trout were repeatedly sampled within lakes in each year, we calculated temporal beta diversity for fish as the mean Bray-Curtis distance across all pairwise fish comparisons between years within each lake. In contrast, bacterioplankton were sampled once per year in each lake, so we calculated temporal turnover as the Bray-Curtis distance between the two annual samples. These yielded lake level estimates of temporal turnover for each habitat type. We modeled differences between habitat types using a GLMM with a beta distribution (logit-link) using habitat type as a fixed effect and lake as a random effect.

Due to differences in sampling intensity between fish and water microbes, we performed additional sensitivity analyses that equalized replication between habitat types. For VI, we repeatedly subsampled fish within each lake without replacement to match the replication of water samples and recomputed differences in within-lake spatial heterogeneity. For the SN, we repeatedly sampled one brook trout per lake-year to match bacterioplankton sample sizes and recalculated differences in within-lake temporal turnover between habitat types. We repeated these procedures for 999 random permutations (Table S9).

### 2.6 Spatial turnover

We estimated the strength of spatial turnover (distance-decay relationships) across the four sample categories using both taxonomic (1 – Bray-Curtis) and phylogenetic (1 – Weighted-Unifrac) measures of community similarity. We filtered out within-lake comparisons and fit linear mixed effects models that predict pairwise community similarity as a function of geographic distance with comparisons from the same year as a random effect using the package glmmTMB (Brooks et al. 2017). The strength of relationships were quantified from model coefficients and marginal R^2^ values were estimated using *r.squaredGLMM()* in the ‘MuMIn’ package (Brooks & Brooks 2015). To examine the statistical significance and correlations of distance-decay relationships, we used Mantel-tests (Spearman, 999 permutations) modified for use on non-square matrices.

### 2.7 Comparing Species Distributions to a Neutral Model

To test whether host-associated and free-living microbes vary in their consistency with neutral expectations, we fit the Sloan Prokaryote Neutral Model (PNM; Sloan et al. 2006) separately to each sample category (brook trout gut, stickleback gut, SN bacterioplankton, VI bacterioplankton) using code adapted from Burns et al. 2016. This stochastic model relates the mean relative abundance of each microbial taxon (ASV) to its frequency of occurrence across local communities under neutral dynamics (random immigration, reproduction, and death). We compared model fit (R^2^) and estimated immigration parameters (m) among sample categories to assess differences in the relative importance of neutral processes. ASVs were classified as occurring “above”, “within”, or “below” neutral model predictions based on confidence intervals around the fitted relationship.

We examined whether these neutral model partitions were phylogenetically structured consistently across habitat types and study systems by constructing presence-absence communities for each combination of sample category and neutral partition. Each of these communities (12 total) represented the set of ASVs classified as above, below, or within the neutral model prediction for a given sample category. We calculated unweighted Unifrac distances (presence-absence) and used PERMANOVA to test for phylogenetic differences among these communities. Neutral model partition habitat type, and their interaction were fixed effects in the model, and we restricted permutations by system to account for sampling differences between the SN and VI. Finally, we tested whether taxa shared between fish and water samples across systems exhibited consistent deviations from neutral expectations by correlating (Spearman) their model deviance values.

## 3. RESULTS

### 3.1 Fish and lake water microbiomes exhibit consistent differences across systems

Independent analysis of each system indicated that communities partition between habitat types (fish gut and bacterioplankton; Fig 2A; Fig 2B; Tables S2-S5). In both the temporal and spatial PERMANOVA models, habitat types were significantly different based on Bray-Curtis (SN: p = 0.001; VI: p = 0.001) and weighted Unifrac (SN: p = 0.001; VI: p = 0.001) distances. Habitat type explained the same amount of community variation in both models for each system, and it consistently explained more variance in the SN than on VI (Bray-Curtis SN: R^2^ = 0.21; weighted Unifrac SN: R^2^ = 0.39; Bray-Curtis VI: R^2^ = 0.09; weighted Unifrac VI: R^2^ = 0.11).

**Figure 1.**
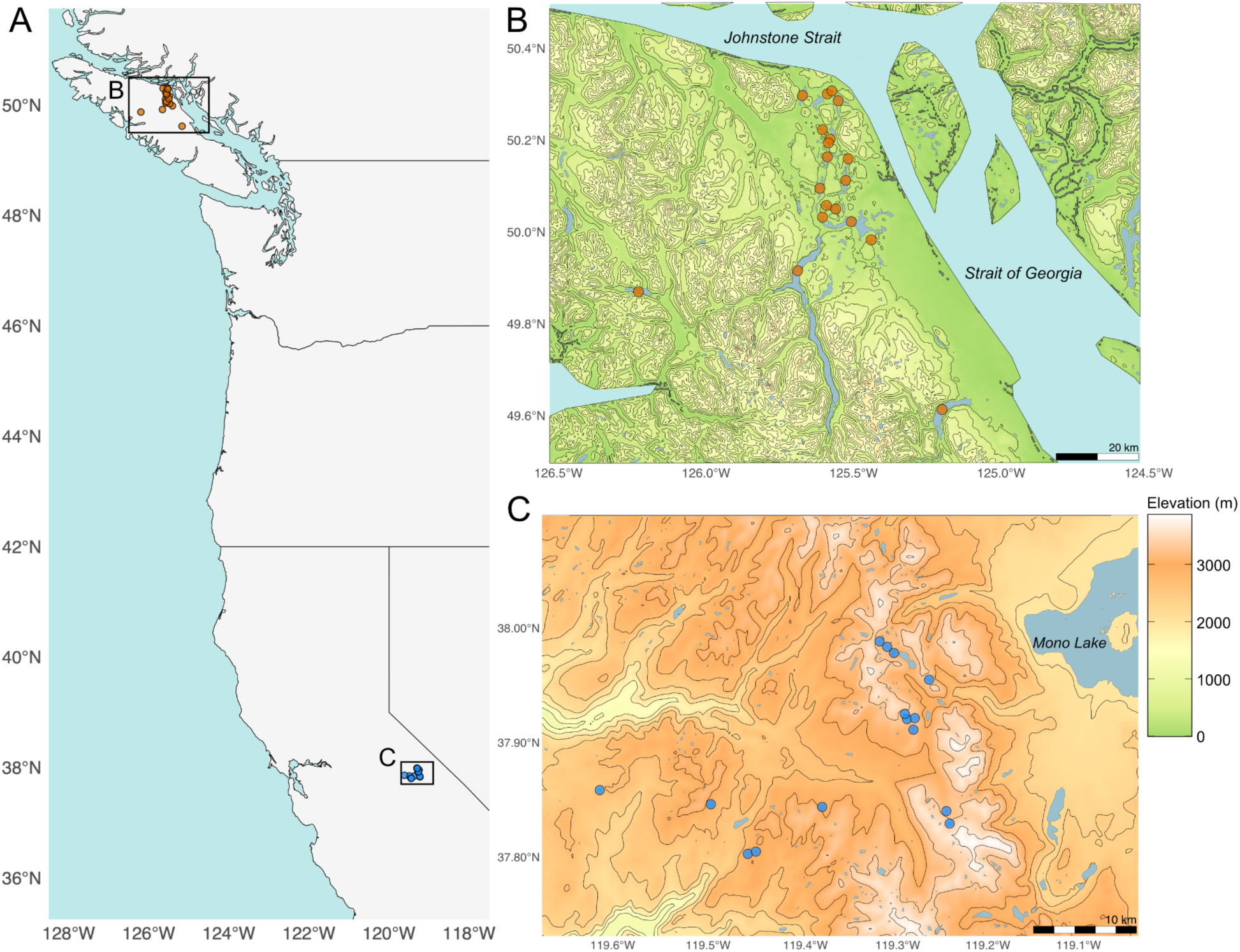
Map of study systems and sites. **(A)** Western portion of North America, boxes represent the locations of study systems. Panels **(B)** and **(C)** represent Vancouver Island and Sierra Nevada respectively. Orange points represent lakes sampled on Vancouver Island. Blue points are lakes sampled in the Sierra Nevada. Contour lines and colors are varying elevations ranging from sea level to 4000m.

**Figure 2.**
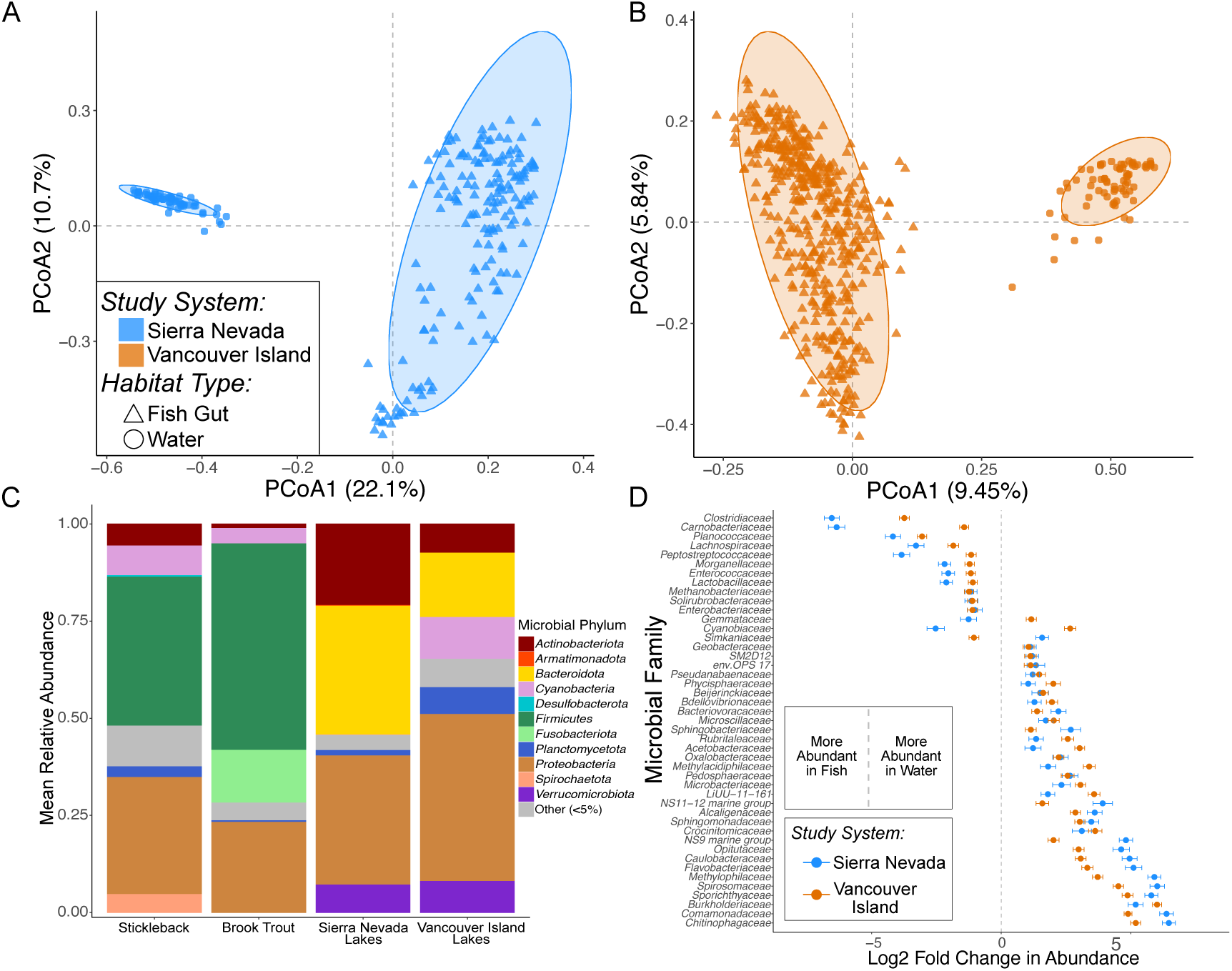
Panels. **(A)** and **B** depict Principal Coordinate Analysis (PCoA) on Bray-Curtis distance matrices for the Sierra Nevada and Vancouver Island study systems respectively. Orange points represent samples collected from Vancouver Island and blue points are samples from the Sierra Nevada. Shapes denote whether microbial communities are from fish guts (*triangles*) or lake water (*circles*). Ellipses represent 95% confidence intervals around the centroid of habitat types (fish gut or water). **(C)** Mean relative abundance barplots of major phyla (>5%) across sample categories. **(D)** Analysis of Compositions of Microbiomes with Bias Correction (ANCOM-BC) depicting the differential abundance of microbial families between fish and lake microbiomes within each system. Comparisons were made independently for each system (Sierra Nevada bacterioplankton vs brook trout; Vancouver Island bacterioplankton vs stickleback). Negative log-fold values indicate families differentially abundant in fish guts and positive log-fold values represent families differentially abundant in water microbiomes.

In the spatial models, lake structured microbiomes in both systems (Tables S2 & S3). However, lake explained less variance than habitat type in the SN for both distance metrics (Bray-Curtis SN: R^2^ = 0.16, p = 0.001; weighted Unifrac SN: R^2^ = 0.10, p = 0.001). In contrast, lake explained a similar amount of variation as habitat type on VI (Bray-Curtis VI: R^2^ = 0.11, p = 0.001; weighted Unifrac VI: R^2^ = 0.11, p = 0.001). Lake by habitat type interaction terms were significant in both systems indicating that space differentially structures host and free-living microbes (p = 0.001 in all models).

In the temporal models, sampling year was significant in both systems but explained less variation than habitat type or lake (Tables S4 & S5; Bray-Curtis SN: R^2^ = 0.04, p = 0.001; Weighted-Unifrac SN: R^2^ = 0.03, p = 0.002; Bray-Curtis VI: R^2^ = 0.03, p = 0.001; Weighted-Unifrac VI: R^2^ = 0.03, p = 0.001). Interaction terms between habitat type and year were significant but explained less variance than the interaction with lake (Bray-Curtis SN: R^2^ = 0.007, p = 0.018; Weighted-Unifrac SN: R^2^ = 0.009, p = 0.012; Bray-Curtis VI: R^2^ = 0.01, p = 0.001; Weighted-Unifrac VI: R^2^ = 0.01, p = 0.001).

The balanced subsampling sensitivity analysis yielded effect sizes and significance patterns consistent with the full models for habitat type, spatial, and temporal effects on VI (Table S8). In the SN, habitat type effects remained strong and significant across all subsampled datasets, confirming that differences between fish and water microbiomes were robust to unequal sample sizes. However, lake and temporal effects were reduced in the SN balanced analyses because fish were not sampled in all years, whereas water microbiomes were sampled across additional years. Consequently, balanced subsampling was restricted to site-year combinations where both sample types were present, reducing spatial and temporal replication. In summary, habitat type remained the dominant and most consistent predictor of microbiome structure across all analyses despite unequal sample sizes (Table S8).

At a course phylogenetic resolution (Phylum), fish gut and water microbiomes show consistent differences across systems (Fig 2C). Fish gut microbiomes were enriched in *Firmicutes* and *Fusobacteriota*, whereas bacterioplankton were dominated by *Bacteroidota* and *Verrucomicrobiota*. At a finer taxonomic resolution, differential abundance analysis identified 79 (SN) and 75 (VI) microbial families that differed between fish guts and bacterioplankton (Fig 2D). Of these differentiated taxa, 45 families were shared across the two systems with only three showing opposite directions of differentiation between systems (Fig 2D). Families consistently enriched in fish included *Clostridiaceae*, *Carnobacteriaceae,* and *Lactobacillaceae*, whereas bacterioplankton were enriched in families such as *Chitinophagaceae*, *Burkholderiaceae*, and *Comamonadaceae*. Overall, these results indicate that fish gut and bacterioplankton microbiomes show reproducible taxonomic contrasts across two independent lake systems.

### 3.2 Alpha and beta diversity of fish gut and bacterioplankton

We found that bacterioplankton had consistently higher alpha diversity than fish gut microbiomes in both systems (Fig 3A; Fig 3B). Fish gut microbiomes exhibited significantly lower ASV richness than bacterioplankton in both systems (Table S6; GLMMs; SN: p < 0.001 VI: p < 0.001). Similarly, Shannon diversity was lower in fish guts compared to bacterioplankton in both the SN (Table S6; Gaussian LMM; SN: p < 2×10⁻^16^) and on VI (Table S6; Gaussian LMM; VI: p < 2×10⁻^16^).

**Figure 3.**
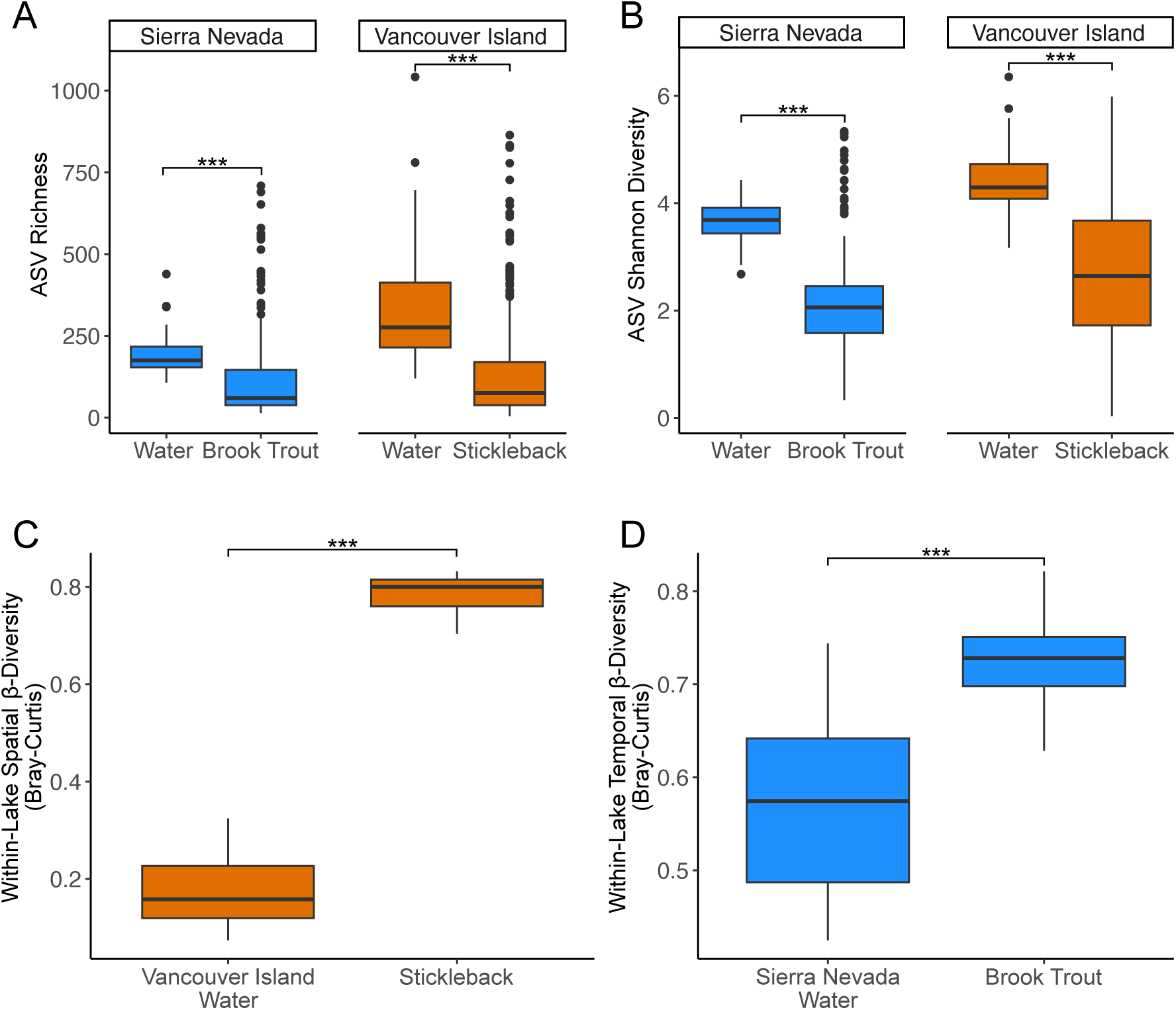
Patterns of contrasting alpha and beta diversity between fish guts and bacterioplankton. Panels **(A)** and **(B)** depict ASV Richness and Shannon Diversity for habitat types (fish gut and water microbes) in each system respectively. **(C)** Within-lake spatial beta diversity for Vancouver Island samples (lake water and stickleback), quantified as the mean Bray-Curtis distance of replicate samples to the lake-level centroid community for each habitat type (mean within-lake heterogeneity). **(D)** Within-lake temporal beta diversity for Sierra Nevada samples (lake water and brook trout), quantified as Bray-Curtis dissimilarity between 2021 and 2022 samples within each lake. For brook trout, values are means across all pairwise fish-fish comparisons between years within a lake. For all panels, asterisks represent significant differences between habitat types (p < 0.05) based on GLMMs.

In contrast, we found consistently higher within-lake beta diversity across space and time in fish gut microbiomes compared to bacterioplankton (Fig 3C; Fig 3D). On VI, mean within-lake spatial beta diversity (all samples collected in the same year) was higher for stickleback gut microbes compared to bacterioplankton samples (Table S7; beta GLMM; p < 2×10⁻^16^). Habitat type explained most of the variation in within-lake beta diversity on VI (marginal R^2^ = 0.99, Table S7) with little additional variance attributable to the random effects of lake or year. In the SN, within-lake temporal beta diversity (comparisons between years in the same lake) was significantly higher in fish gut microbiomes compared to bacterioplankton (Table S7; beta GLMM; p = 4.53×10^-14^) and habitat type explained a majority of the variance (marginal R^2^ = 0.71). Our sensitivity analysis confirmed that these results were not sensitive to differences in sample size between fish and bacterioplankton as 100% of model permutations with equal sample size found higher beta diversity in fish gut microbiomes for both metrics (Table S9). Overall, these results indicate that fish gut microbes are less diverse at the local scale but more differentiated within lakes over space and time.

### 3.3 Spatial turnover

We found that fish gut and lake water microbiomes exhibit spatial turnover (distance-decay) using Bray-Curtis similarity (Fig 4). However, fish exhibited weaker relationships (brook trout Mantel r = –0.16, p = 0.001; stickleback Mantel r = –0.11, p = 0.001) compared to lakes (SN lakes Mantel r = –0.37, p = 0.001; VI lakes Mantel r = –0.36, p = 0.001). Model slopes were also greater in lake microbes (SN lakes β = –3.8e-3; VI lakes β = –1.3e-3) compared to fish guts (brook trout β = –2.4e-3; stickleback β = –2.9e-4). Geographic distance explained more variation in lake microbiomes (SN lakes R^2^ = 0.13; VI lakes R^2^ = 0.10) compared to fish (brook trout R^2^ = 0.03; stickleback R^2^ = 0.008).

**Figure 4.**
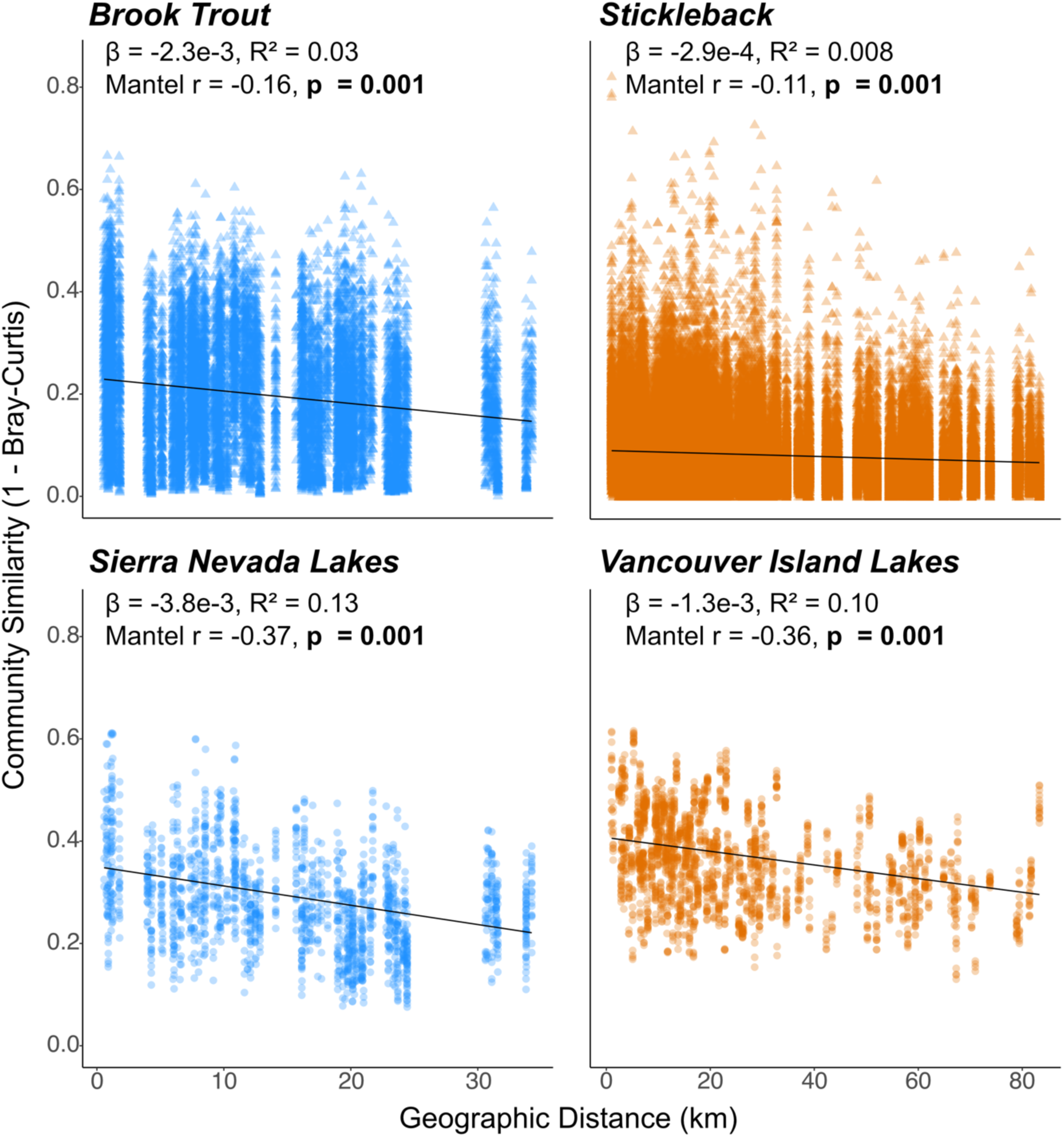
Distance decay relationships depicting pairwise community similarity (1 – Bray-Curtis) as a function of geographic distance between samples for brook trout, stickleback, Sierra Nevada lakes, and Vancouver Island lakes. Points are colored by study system (*orange* – Vancouver Island, *blue* – Sierra Nevada) and shape denotes sample category (*circle* – water, *triangle* – fish gut). Slopes and adjusted R^2^ values were calculated from linear mixed effects models with comparisons for communities from the same year as a random effect. Mantel r values and the corresponding p-values are reported for determining the statistical significance of relationships.

Phylogenetic distance-decay relationships were weaker and less consistent across systems, and geographic distance explained less phylogenetic turnover than taxonomic turnover (Fig S2). Partial Mantel tests indicated that elevation contributed to spatial turnover in Sierra Nevada brook trout microbiomes independent of geographic distance but had weaker or inconsistent effects on lake microbiomes (Table S10). Together, these results indicate that bacterioplankton exhibit stronger and more consistent spatial structuring than host-associated microbiomes.

### 3.4 Neutral model analysis

The prokaryote neutral model explained more variation in bacterioplankton communities (Fig 5; SN bacterioplankton R^2^ = 0.63, VI bacterioplankton R^2^ = 0.74) compared to fish gut microbiomes (Fig 5; brook trout R^2^ = 0.33, stickleback R^2^ = 0.35). Fish guts also contained fewer widely distributed (high-occurrence) ASVs relative to lake water communities (Fig 5). Estimated immigration parameters (m) followed the same pattern: fish metacommunities exhibited lower immigration rates (brook trout m = 0.006; stickleback m = 0.002) than bacterioplankton (SN m = 0.017; VI m = 0.098). Most ASVs fell within neutral expectations (within model prediction) across all sample categories (Fig S3A; brook trout: 75%; stickleback: 84%; SN bacterioplankton: 89%; VI bacterioplankton: 86%). However, the distribution of non-neutral ASVs differed between habitat types. Fish microbiomes had a greater proportion of ASVs occurring more frequently than expected (“above”; brook trout: 24.1%; stickleback: 15.5%) compared to bacterioplankton (SN bacterioplankton: 9.5%; VI bacterioplankton: 11.5%). In contrast, bacterioplankton metacommunities had slightly more ASVs that occur less frequently than expected (“below”; brook trout: 0.9%; stickleback: 0.5%; SN bacterioplankton: 2%; VI bacterioplankton: 3%). Overall, these results indicate stronger departure from neutral expectations in fish gut microbiomes relative to lake bacterioplankton.

**Figure 5.**
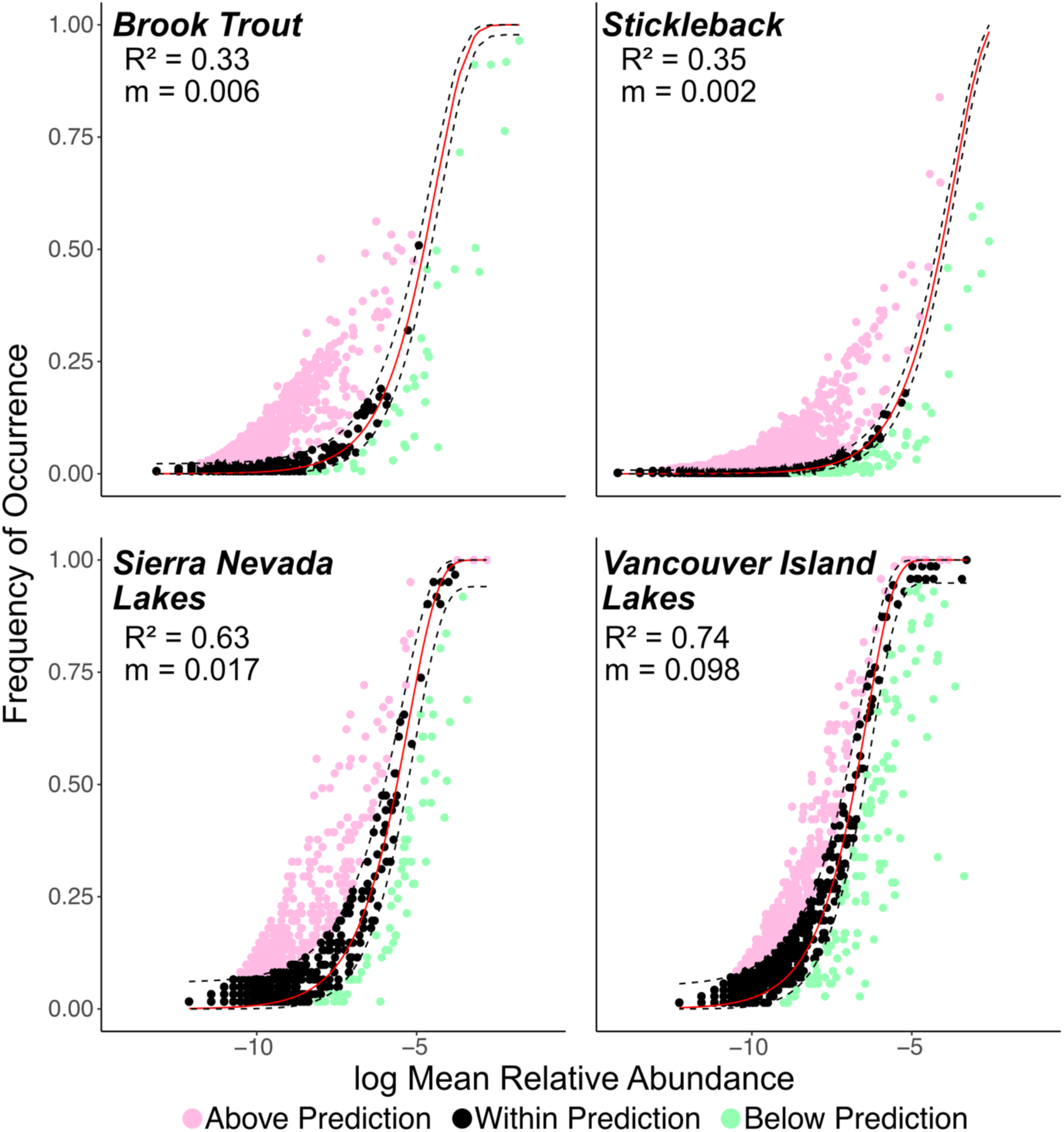
Plotted fit (R^2^) and estimated immigration rate (m) from the Prokaryote Neutral Model (PNM) for brook trout, Sierra Nevada lakes, stickleback, and Vancouver Island lakes. The model predicts the frequency of ASV occurrence across all microbial communities as a function of their respective log mean relative abundance within local communities and assumes neutral dynamics. Solid red and black dashed lines are the model predictions and confidence intervals respectively. Pink, black, and green points represent individual ASVs above, within, and below the confidence interval of the predicted frequency of occurrence

We found phylogenetic differences between the microbes found above, below or within the neutral model predictions (Table S11; Fig S3B). Neutral model partition explained significant variation in community composition based on unweighted Unifrac distances (PERMANOVA; R^2^ = 0.31, p = 0.001). Habitat type also explained variance (R^2^ = 0.14, p = 0.007). However, the interaction between selection partition and habitat type explained more variation than habitat type (selection x habitat type; R^2^ = 0.20, p = 0.028), indicating that phylogenetic differentiation among neutral model partitions differed between fish and bacterioplankton communities. In particular, neutral and above-prediction ASVs were more phylogenetically similar in fish gut microbiomes than in bacterioplankton (Fig S3B). These partitions were associated with distinct taxa that were repeatable between fish and water microbiomes across systems (Fig 6A). In addition to phylogenetic similarities, we found that deviances of ASVs present in fish and lake water from both systems were significantly correlated (Fig 6B; Fig 6C). In bacterioplankton, 87% of ASVs present in both systems exhibited consistent deviation from the model (6B). In fish, 98% of shared ASVs showed consistent deviation (Fig. 6C), suggesting that non-neutral processes may be stronger in fish gut microbiomes (6C).

**Figure 6.**
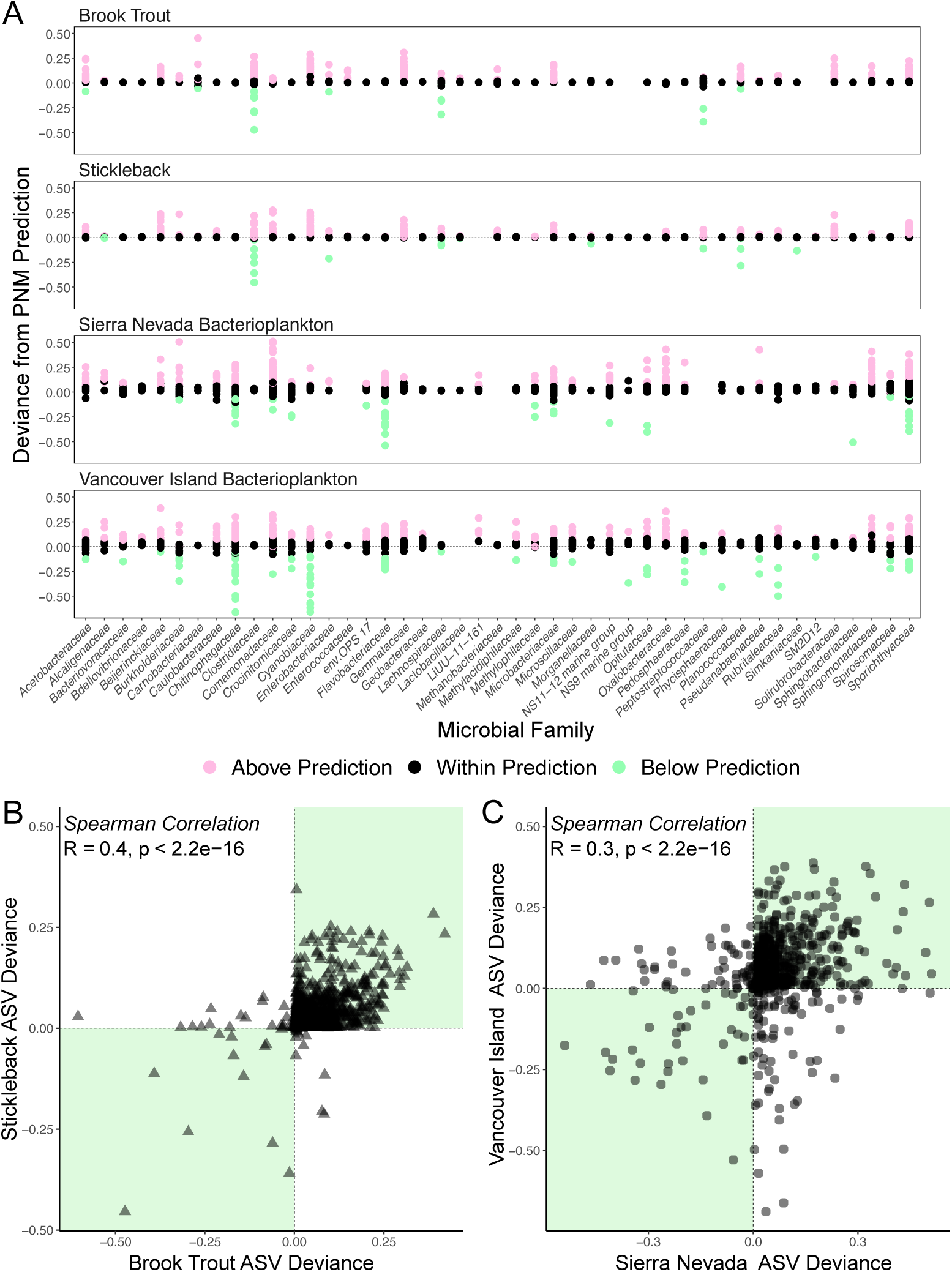
(**A**) Deviance in the predicted frequency of occurrence from the PNM for ASVs across microbial families that were differentiated between habitat types. ASVs are color coded by above (*pink*), within (*black*), and below (*green*) the model confidence interval. Panels **(C)** and **(D)** depict scatterplots of ASV deviances from the model prediction between hosts and lakes respectively. Green quadrants represent regions of repeatability between systems. For fish and water, 98% and 87% of shared ASVs exhibited repeatable deviance from the model prediction respectively.

## 4. DISCUSSION

We found that host-associated and free-living microbial communities from fish guts and lake water have different metacommunity structure and biogeographic patterns across two independent systems. First, bacterioplankton and fish gut microbiomes diverged in community composition across two geographic regions with two fish host species. Second, in both regions, fish microbiomes were less diverse at the local scale but had greater turnover within lakes. This combination suggests reduced exchange of microbes between fish hosts than between lakes, either due to reduced horizontal transmission or stronger host selection constraining colonization. Fish microbiomes had lower spatial turnover than free-living lake microbes. Finally, the neutral model better predicted patterns of abundance and occurrence in free living microbes than fish gut microbiomes, suggesting that non-neutral processes shape host-associated microbiomes more strongly. These differences in biogeography, diversity, and sensitivity to neutral ecological processes between host-associated and free-living microbes suggest fundamental differences in how ecological processes shape microbial communities across environments.

We found consistent variation in diversity between fish gut and lake microbes across study systems. Fish guts had significantly lower alpha diversity compared to water microbes in both ASV richness and Shannon diversity (Fig 3). Conversely, beta diversity within lakes over space and time was greater in fish hosts. The similar patterns across distinct systems suggest the processes driving metacommunity structure vary in magnitude and scale among host and environmental microbes.

One potential explanation for this result may be differences in dispersal. Previous work has demonstrated that dispersal can shape the diversity of microbes in the gut (Burns et al. 2017) and in the environment (Evans et al. 2017; Hawkins & Zeglin 2022; Louca 2022). Theory predicts that slower dispersal through the environment constrains local diversity and elevates beta diversity across habitat patches (Loreau & Mouquet 1999). In fish hosts, association with a discrete and selective habitat patch may limit opportunities for microbial exchange among individuals or populations, potentially contributing to lower alpha diversity and elevated beta diversity within and across lakes. In contrast, free-living microbes may experience more rapid dispersal within and across lakes that may amplify local alpha diversity while depressing beta diversity. Immigration estimates from the prokaryote neutral model are consistent with this interpretation, as fish gut microbes exhibited consistently lower neutral model immigration estimates compared to lake bacterioplankton (Fig 5). While neutral models do not directly quantify dispersal, the lower inferred migration rates suggest potentially stronger dispersal limitation in hosts relative to free-living microbes. Hosts may constrain the movement of their associated microbes within and across habitats, depressing local diversity and elevating turnover across space and time. In contrast, higher immigration estimates for free-living microbes align with lower beta diversity within lakes among bacterioplankton samples. Similar patterns in bacterioplankton have been observed in Sierra Nevada alpine lakes (Wall & Perreault et al. 2025; Härer et al. 2025), supporting the view that free-living microbial communities are comparatively well mixed relative to fish microbiomes. Together, these results are consistent with differences in dispersal between host-associated and environmental microbes, though direct measures of microbial dispersal and selection among habitat patches are required to confirm this mechanism.

Differences in diversity patterns may also reflect variation in the magnitude and scale of ecological selection between host-associated and free-living microbes. As habitat heterogeneity increases, the importance of species-sorting should increase, driving higher beta diversity and constraining alpha diversity due to tight environmental filtering. Fish hosts may represent more heterogeneous and selective habitats than the surrounding water column, given inter-individual differences in genotype, physiology, and diet. Individual specialization and dietary variation may be particularly important in shaping local and beta diversity patterns in fish hosts as diet diversity and composition shape gut microbiomes (Bolnick et al. 2014). In stickleback and brook trout, there is substantial within and across population variation in niche width and diet diversity associated with environmental gradients (Bolnick et al. 2020; Baker et al. 2022). Individual specialization for specific diet items may alter the selection of microbial specialists that metabolize diet compounds (Ingala et al. 2021) which may scale up to affect host foraging behavior (Trevelline & Kohl 2022). In free-living lake microbes, ecological selection likely operates on temporal and spatial scales associated with external environmental heterogeneity. For example, microbial alpha and beta diversity is associated with environmental conditions related to climate and nutrients in mountain lakes (Schulhof et al. 2020), but these forces may be more spatially homogenized within lakes (Härer et al. 2025) than host-associated filters are among individuals. Our results suggest that potentially stronger ecological selection in hosts compared to environmental microbes may drive disparate patterns of local, regional, and turnover components of diversity.

We found differences in spatial turnover (distance-decay) between fish and lake water microbiomes. In fish microbiomes we found statistically significant, although weak, distance-decay patterns using Bray-Curtis similarity across both systems (Fig 4). These patterns were highly variable, demonstrating that some hosts exhibit differentiated microbiomes even across small spatial scales that qualitatively align with within-lake beta diversity estimates (Fig 3; Fig 4). In contrast, bacterioplankton displayed uniformly stronger distance-decay relationships with less variation (Fig 4), suggesting that turnover (beta diversity) is connected to spatial and environmental gradients. Our observations mirror past work demonstrating that fish microbiomes exhibit weaker distance-decay patterns relative to their free-living counterparts, and that external environmental conditions structured free-living communities but explained little variation in fish guts (Suzzi et al. 2023). These results suggest that fish gut microbiomes are primarily structured by host-level ecological selection and dispersal limitation, resulting in weak spatial structuring across sites. However, elevation was associated with variation in brook trout gut microbiome composition, indicating that climate gradients can influence host-associated microbial communities. This result connects to prior work demonstrating connections between climate and brook trout ecology (Symons & Shurin 2016; Symons et al. 2019). In contrast, free-living microbes in the water column had stronger spatial turnover, suggesting that communities are assembled at geographic, landscape level scales; this pattern may be driven by factors such as watershed hydrology/connectivity, wind, environmental heterogeneity, and physical barriers to dispersal (Crump et al. 2007; Bottos et al. 2020; Maltz et al. 2022). Our data suggest that the scale at which ecological processes operate differs between host-associated and environmental microbiomes, producing distinct patterns of spatial turnover.

The neutral model better described abundance-occurrence relationships in lake microbes than in fish guts (Fig 5). The neutral model explained more than half of the variance in lake microbes but less than forty percent across both fish hosts (Fig 5). While most taxa were classified as “neutral” across all sample categories, a higher proportion were non-neutral in fish, particularly in brook trout (Fig S3). These patterns suggest that the abundance-occurrence relationships of microbes in fish guts conform less to neutral expectations. Prior work has shown that stochastic processes explain the structure of microbial communities in both environmental (Sloan et al. 2006; Woodcock et al. 2007; Ofiţeru et al. 2010) and host-associated (Heys et al. 2020; Henry & Ayroles 2022) samples. However, the reduced model fit and higher proportion of non-neutral ASVs in fish hosts suggest a potentially larger role for processes not captured by the neutral model including host selection and non-neutral dispersal limitation. These results connect with our observed patterns of diversity and spatial turnover where hosts exhibit constrained local diversity and higher overall turnover consistent with potentially stronger deterministic forces. Conversely, lake bacterioplankton more clearly adhere to neutral expectations, suggesting that stochastic processes may explain more variance in the relationship between abundance and occurrence for microbes in lakes.

We found that fish gut metacommunities contained a higher proportion of taxa classified as “above” the model prediction (Fig S4) that are widely distributed but relatively rare. These may represent taxa that are under consistent environmental filtering or possess traits the promote dispersal/colonization across hosts (Echenique-Subiabre et al. 2025). However, their generally low relative abundances within local communities suggest that host selection may also limit their abundance. One possible explanation is that taxa with strong dispersal or colonization abilities may experience competition tradeoffs that constrain their abundance in local habitat patches (Dini-Andreote et al. 2018; Smith et al. 2018; Wetherington et al. 2022). In contrast, bacterioplankton contained fewer “above” ASVs, demonstrating that processes such as strong environmental filtering and non-neutral dispersal may be weaker in these communities.

Model partitions formed distinct phylogenetic groups that were similar between the Sierra Nevada and Vancouver Island (Fig 6A; Fig S4B). Taxa shared across systems between fish hosts and lakes exhibited similar deviances from the model prediction (Figs 6C and 6D). This combination suggests that closely related lineages tend to fall into similar abundance-occurrence categories. However, the degree of separation between neutral and non-neutral taxa varied between fish guts and lake water samples (Fig S4B). Specifically, neutral and “above” microbes were more phylogenetically similar in fish guts compared to bacterioplankton samples. Our results emphasize the utility of neutral models for identifying taxa that may be consistently structured by host-associated and environmental factors. Further studies should analyze the functional mechanisms underlying differences in the abundance-occurrence patterns of neutral and non-neutral microbes to more clearly link patterns to processes. Those that fall above the model prediction may exhibit traits associated with higher dispersal capabilities or key metabolic functions central to host physiology or ecosystem functioning. In contrast, taxa falling below the model prediction could potentially be pathogenic/parasitic for hosts or may be mismatched to external environmental conditions. Overall, we show that microbes in fish guts and lakes exhibit different sensitivity to neutral processes and the same taxa often deviate from abundance-occurrence predictions across two distinct study systems.

Our study has implications for understanding the distribution of microbial diversity across hosts, environments, space, and time. The data indicate that host-associated (fish gut) and free-living (lake bacterioplankton) microbiomes exhibit distinct patterns of diversity, spatial turnover, and consistency with neutral abundance-occurrence relationships. These differences are consistent with ecological processes such as dispersal and species-sorting operating at different spatial and biological scales in host-associated versus free-living habitats. Critically, our results were consistent across two case studies with different ecosystems, hosts, and sampling designs. This suggests that host-association may fundamentally alter the distribution and diversity of microbes across ecosystems. Future work should test the generality of these patterns across hosts, environments, and spatial scales to uncover the processes driving microbial community assembly and their effects on hosts and ecosystems.

## ACKNOWLEDGEMENTS

We thank director Carol Blanchette and the Sierra Nevada Research Laboratory for lodging, laboratory space, and logistical support. Thank you to Henry K. Baker for assistance in study design and sampling. This project was supported by multiple awards including the NSF GRFP (to JHD), UCSD Jeanne M. Messier Memorial Award (to JHD), and UC Natural Reserve System Mildred E. Mathias Graduate Student Research Grant (to JHD). Additional funding was provided by NSF (award number 2018058).

## DATA ACCESSIBILITY AND BENEFITS-SHARING

Code and raw data are available on Dryad (reviewer link: http://datadryad.org/share/IdA6BauwDU22YTkWnpj5Z3VK2K2Yc-KkU58ptrZ3JTQ). Sequencing data were submitted to NCBI SRA at Project PRJNA1298366. Benefits Generated: Samples used in this study were collected in the United States and Canada in compliance with institutional permits and national regulations. No additional access and benefit-sharing obligations apply.

## AUTHOR CONTRIBUTIONS

JHD, AH, DJR, and JBS conceived of and designed the study; JHD, AH, CBW, CCS collected and processed data; JHD and AH analyzed the data; JHD wrote the manuscript; AH, CBW, CCS, DJR, and JBS revised the manuscript; JHD, JBS, and DJR acquired funding.

