## Supporting Information for "Consistent, scale-dependent differences in the biogeography of host-associated and free-living microbiomes across systems"

Joshua H. Dominguez (JHD)^1^

Andreas Härer (AH)^1^

Christopher B. Wall (CBW)^1,2^

Diana J. Rennison (DJR)^1^

Celia C. Symons (CCS)^3^

Jonathan B. Shurin (JBS)^1^

**^1^** School of Biological Sciences, Department of Ecology, Behavior, & Evolution, University of California San Diego, La Jolla, California, USA

**^2^** Department of Earth Sciences, University of Hawai‘i at Mānoa, Honolulu, Hawaii, USA

^3^ Department of Ecology, Behavior and Evolution, University of California, Irvine, California, USA

**METHODS 1 – Sampling Protocols**

We sampled free-living microbes from lake water in 15 Sierra Nevada lakes during the summer seasons of 2016, 2017, 2019, 2021, and 2022. Using an inflatable kayak, we located the deepest point in each lake using a depth sounder and collected water from 1m below the surface using an integrated vertical tube sampler. Water was pre-filtered through a 64-μm mesh to sift out large particles and between 300 and 1080 mL of water was hand filtered through a bleach-sterilized filter apparatus onto a 0.22 μm Sterivex filter cartridge (Millipore) and kept on ice in the field. Once back at the lab, we stored Sterivex filters at -80 °C for downstream processing. Across the five sampling years we collected a total of 63 lake water samples (n = 2 – 5 samples per lake). In 2021 and 2022 we collected fecal material from 184 brook trout. Fish were captured via angling using barbless lures/flies and fecal material was collected using a non-lethal protocol. After capture, individuals were inverted exposing the ventral side, and fecal material was expelled from the anal cavity onto sterile cotton swabs by sliding a thumb with light pressure from the cranial to the caudal end along the ventral side of the animal. We then released individuals after allowing for adequate recovery. Cotton swabs with fecal material were placed in sterile 2 mL microcentrifuge tubes and kept on ice in the field before final storage at -80°C in the laboratory.

We sampled free-living water microbes by hand-filtering 120 mL of lake water collected from the shore using a bleach sterilized syringe equipped with a 0.2 µm filter (Whatman plc, Maidstone, UK). At each lake we collected 3 – 4 replicate filters totaling 74 water samples. After collection, filters were stored at -20°C until DNA extraction. We captured 541 stickleback (n = 16 – 30 individuals per lake) using minnow traps under British Columbia collection permits NA20-602264, MRVI21-619908, and NA22-713085. Fish were sacrificed using a lethal dose of MS-222 (500 mg/L) and stored at -20°C. We removed whole intestines from individuals using sterile dissection tools and carefully excreted gut contents by gentle squeezing. After dissection, guts were stored in sterile 2 mL microcentrifuge tubes at -80°C for downstream DNA extraction

**METHODS 2 – Library Preparation**

We amplified the V4 Region of the 16S rRNA gene using 515F-806R primer sets with Illumina adapters for all samples (Caparosa et al. 2012). For Sierra Nevada samples we used modified primer sets that increase detection of aquatic bacteria (Apprill et al. 2015; Parada et al. 2016; Walters et al. 2016).

Genomic DNA from Sierra Nevada lake filters was extracted using the Qiagen DNEASY PowerWater kit and sent samples to the Argonne National Laboratory for downstream library preparation. PCR mixes consisted of 9.5 µL of MO BIO PCR Water (Certified DNA-Free), 12.5 µL of QuantaBio’s AccuStart II PCR ToughMix (2x concentration), 1 µL Golay barcode tagged Forward Primer (5 µM concentration), 1 µL Reverse Primer (5 µM concentration), and 1 µL of template DNA. Thermocycler conditions consisted of 94 °C for 3 minutes, 35 cycles at 94 °C for 45 s, 50 °C for 60 s, 72 °C for 90 s; and 72 °C for 10 minutes. Amplicons are then quantified using PicoGreen (Invitrogen). Once quantified, volumes of each of the products were pooled into a single tube. The pool was cleaned using AMPure XP Beads (Beckman Coulter) and quantified using a fluorometer (Qubit, Invitrogen). Amplicons were sequenced on a PE250 MiSeq run.

Genomic DNA was extracted from Brook Trout fecal material using the Qiagen PowerSoil kit according to manufacturer protocols and a two-step PCR reaction was performed. PCR mixes and conditions were modified to optimize for brook trout samples and consisted of 5 uL of Forward Primer (1x concentration), 5 uL of Reverse Primer (1x concentration), 12.5 uL of 2X KAPA HiFi HS ReadyMix, and 2.5 uL of DNA. Thermocycler conditions were 95 °C for 3 minutes, 28 cycles of 95 °C for 30 s, 55 °C for 30 s, 72 °C for 30 s; and 72 °C for 5 minutes. Successful amplification was visualized using gel electrophoresis (2% agarose gel). A second round of PCR was performed to attach Illumina barcodes and consisted of 5 uL of i7 primer (5 nm), 5 uL of i5 primer (5 nm), 25 uL of 2X KAPA HiFi HS ReadyMix, 10 uL of DPEC Water, and 5 uL of round 1 PCR product. Amplified, barcoded DNA was quantified, cleaned using AMPure XP beads, pooled, and quality checked with a BioAnalyzer. Sequencing was performed on a PE250 MiSeq run at the University of California –Davis GenomeCenter.

Genomic DNA from Vancouver Island samples (stickleback gut and lake water) was extracted using the Qiagen PowerSoil kit. PCR was performed in triplicate using a 10 uL reaction volume with the Platinum II Hot Start PCR master mix (Thermo Fisher Scientific, Waltham, MA), and the three replicates were subsequently pooled. Thermocycler conditions were 98 °C for 60 s, 35 cycles of 98 °C for 10 s, 56 °C for 20 s, 72 °C for 60 s; and an elongation of 72 °C for 10 minutes. Gel electrophoresis (2% agarose) was performed to check for amplification. DNA concentrations were measured on a Qubit 4 fluorometer (Thermo Fisher Scientific), samples were pooled in an equimolar manner, bead cleaned, and quality checked with a bioanalyzer. Libraries were sequenced on the Illumina MiSeq 600 (PE300) platform at the University of California–Davis GenomeCenter.

**TABLES**

**Table S1.** Number of fish and water samples collected from each lake on Vancouver Island and in the Sierra Nevada.

| ***Lake*** | ***System*** | ***No. fish*** | ***No. water samples*** | ***Latitude*** | ***Longitude*** |
| --- | --- | --- | --- | --- | --- |
| Cascade | Sierra Nevada | 9 | 4 | 37.989774 | -119.30626 |
| Elizabeth | Sierra Nevada | 9 | 4 | 37.845185 | -119.36991 |
| Gardisky | Sierra Nevada | 10 | 5 | 37.956277 | -119.25151 |
| Granite 1 | Sierra Nevada | 14 | 3 | 37.9217949 | -119.27583 |
| Granite 2 | Sierra Nevada | 10 | 2 | 37.926453 | -119.27843 |
| Greenstone | Sierra Nevada | 18 | 4 | 37.979647 | -119.29034 |
| Helen | Sierra Nevada | 12 | 4 | 37.830661 | -119.22873 |
| Gaylor 1 | Sierra Nevada | 16 | 5 | 37.9126276 | -119.26885 |
| Gaylor 2 | Sierra Nevada | 16 | 5 | 37.9226947 | -119.26738 |
| May | Sierra Nevada | 6 | 5 | 37.847477 | -119.49308 |
| Spillway | Sierra Nevada | 15 | 5 | 37.841521 | -119.23244 |
| Sunrise 1 | Sierra Nevada | 5 | 5 | 37.804151 | -119.45218 |
| Sunrise 3 | Sierra Nevada | 13 | 4 | 37.806216 | -119.44326 |
| Wasco | Sierra Nevada | 10 | 2 | 37.985032 | -119.29792 |
| Lukens | Sierra Nevada | 6 | 5 | 37.85989 | -119.6161 |
| Amor | Vancouver Island | 18 | 4 | 50.167528 | -125.559361 |
| Bob | Vancouver Island | 23 | 3 | 50.304472 | -125.559361 |
| Boot | Vancouver Island | 30 | 4 | 50.053889 | -125.530917 |
| Brewster | Vancouver Island | 29 | 4 | 50.098806 | -125.584556 |
| Comox | Vancouver Island | 21 | 4 | 49.617 | -125.170833 |
| Echo | Vancouver Island | 26 | 4 | 49.986417 | -125.411 |
| Lawson | Vancouver Island | 29 | 4 | 50.035972 | -125.575889 |
| Lil Goose | Vancouver Island | 25 | 4 | 50.163139 | -125.488778 |
| Lil Mud | Vancouver Island | 29 | 4 | 50.206111 | -125.550361 |
| Lower Campbell | Vancouver Island | 30 | 4 | 50.026167 | -125.478389 |
| Lower Stella | Vancouver Island | 18 | 4 | 50.310944 | -125.544028 |
| Mccreight | Vancouver Island | 28 | 4 | 50.300944 | -125.6435 |
| McNair | Vancouver Island | 26 | 4 | 50.226444 | -125.576389 |
| Merrill | Vancouver Island | 28 | 4 | 50.061444 | -125.562861 |
| Mohun | Vancouver Island | 27 | 4 | 50.116194 | -125.497361 |
| Muchalat | Vancouver Island | 23 | 4 | 49.873528 | -126.199389 |
| Mud | Vancouver Island | 30 | 4 | 50.197639 | -125.555889 |
| Stella | Vancouver Island | 28 | 4 | 50.289389 | -125.522722 |
| Upper Campbell | Vancouver Island | 19 | 4 | 49.919389 | -125.660278 |

**Table S2.** Spatial PERMANOVA (strata = year) output tables for Sierra Nevada samples.

| ***Bray-Curtis*** | *df* | *SS* | *R^2^* | *F* | *Pr(>F)* |
| --- | --- | --- | --- | --- | --- |
| Habitat Type | 1 | 18.4 | 0.21 | 79.3 | **0.001** |
| Lake | 14 | 14.2 | 0.16 | 4.37 | **0.001** |
| Habitat Type * Lake | 14 | 9.10 | 0.10 | 2.81 | **0.001** |
| Residual | 200 | 46.3 | 0.53 |  |  |
| Total | 229 | 88.0 | 1.00 |  |  |
| ***Weighted-Unifrac*** |  |  |  |  |  |
| Habitat Type | 1 | 1.61 | 0.39 | 178 | **0.001** |
| Lake | 14 | 0.42 | 0.10 | 3.35 | **0.001** |
| Habitat Type * Lake | 14 | 0.28 | 0.07 | 2.19 | **0.001** |
| Residual | 200 | 1.80 | 0.44 |  |  |
| Total | 229 | 4.10 | 1.00 |  |  |

PERMANOVA tables generated from 999 permutations; *df = degrees of freedom; SS = sum of squares. Significant effects (p < 0.05) are in bold.*

**Table S3**. Spatial PERMANOVA (strata = year) output tables for Vancouver Island samples.

| ***Bray-Curtis*** | *df* | *SS* | *R^2^* | *F* | *Pr(>F)* |
| --- | --- | --- | --- | --- | --- |
| Habitat Type | 1 | 21.1 | 0.09 | 60.9 | **0.001** |
| Lake | 18 | 26.3 | 0.11 | 4.21 | **0.001** |
| Habitat Type * Lake | 18 | 13.0 | 0.05 | 2.08 | **0.001** |
| Residual | 520 | 180 | 0.75 |  |  |
| Total | 557 | 240.8 | 1.00 |  |  |
| ***Weighted-Unifrac*** |  |  |  |  |  |
| Habitat Type | 1 | 1.32 | 0.11 | 74.1 | **0.001** |
| Lake | 18 | 1.33 | 0.11 | 4.15 | **0.001** |
| Habitat Type * Lake | 18 | 0.62 | 0.05 | 1.95 | **0.001** |
| Residual | 520 | 9.25 | 0.74 |  |  |
| Total | 557 | 12.5 | 1.00 |  |  |

PERMANOVA tables generated from 999 permutations; *df = degrees of freedom; SS = sum of squares. Significant effects (p < 0.05) are in bold.*

**Table S4.** Temporal PERMANOVA (strata = lake) output tables for Sierra Nevada samples.

| ***Bray-Curtis*** | *df* | *SS* | *R^2^* | *F* | *Pr(>F)* |
| --- | --- | --- | --- | --- | --- |
| Habitat Type | 1 | 18.4 | 0.21 | 62.2 | **0.001** |
| Year | 4 | 3.10 | 0.04 | 2.63 | **0.001** |
| Habitat Type * Year | 1 | 0.66 | 0.007 | 2.22 | **0.018** |
| Residual | 223 | 65.9 | 0.75 |  |  |
| Total | 229 | 88.0 | 1.00 |  |  |
| ***Weighted-Unifrac*** |  |  |  |  |  |
| Habitat Type | 1 | 1.61 | 0.39 | 153 | **0.001** |
| Year | 4 | 0.11 | 0.03 | 2.66 | **0.002** |
| Habitat Type * Year | 1 | 0.04 | 0.009 | 3.78 | **0.012** |
| Residual | 223 | 2.35 | 0.57 |  |  |
| Total | 229 | 4.1 | 1.00 |  |  |

PERMANOVA tables generated from 999 permutations; *df = degrees of freedom; SS = sum of squares. Significant effects (p < 0.05) are in bold.*

**Table S5**. Temporal PERMANOVA (strata = lake) output tables for Vancouver Island samples.

| ***Bray-Curtis*** | *df* | *SS* | *R^2^* | *F* | *Pr(>F)* |
| --- | --- | --- | --- | --- | --- |
| Sample Type | 1 | 21.1 | 0.09 | 55.8 | **0.001** |
| Year | 2 | 7.42 | 0.03 | 9.79 | **0.001** |
| Sample Type * Year | 2 | 2.92 | 0.01 | 3.85 | **0.001** |
| Residual | 552 | 209.3 | 0.87 |  |  |
| Total | 557 | 240.8 | 1.00 |  |  |
| ***Weighted-Unifrac*** |  |  |  |  |  |
| Sample Type | 1 | 1.32 | 0.11 | 68.3 | **0.001** |
| Year | 2 | 0.38 | 0.03 | 9.79 | **0.001** |
| Sample Type * Year | 2 | 0.18 | 0.01 | 4.54 | **0.001** |
| Residual | 552 | 10.6 | 0.85 |  |  |
| Total | 557 | 12.5 | 1.00 |  |  |

PERMANOVA tables generated from 999 permutations; *df = degrees of freedom; SS = sum of squares. Significant effects (p < 0.05) are in bold.*

**Table S6.** Generalized Linear Mixed Effects Models Predicting Alpha Diversity in each system.

|  | *Habitat Type Effect Size* | | *Std. Error* | *z-value* | *Pr(>\|z\|)* | *R_m_^2^* |
| --- | --- | --- | --- | --- | --- | --- |
| ***Sierra Nevada*** | |  |  |  |  |  |
| ASV Richness | | 0.76 | 0.12 | 6.34 | **2.35x10^-10^** | 0.16 |
| Shannon Diversity | | 1.53 | 0.13 | 11.9 | **< 2 x10^-16^** | 0.37 |
| ***Vancouver Island*** | |  |  |  |  |  |
| ASV Richness | | 0.98 | 0.11 | 9.04 | **< 2 x10^-16^** | 0.15 |
| Shannon Diversity | | 1.73 | 0.16 | 11.1 | **< 2 x10^-16^** | 0.18 |

Generalized Linear Mixed Effects Models (GLMMs) predicting ASV Richness and Shannon Diversity as a function of habitat type (fish or bacterioplankton) in each system. Richness models were fit with a negative binomial distribution with a log link and Shannon diversity models were fit with a Gaussian distribution. In the Sierra Nevada, lake and year nested within lake were random intercepts. On Vancouver Island, lake and year were independent random intercepts. Columns indicate the system, effect size of habitat type (log space for ASV richness), the standard error of the effect size estimate, z-value, P-value (significant effects are bold), and the marginal R^2^ of models which represents the amount of variation explained by habitat type. Lognormal marginal R^2^ values were used for negative binomial models*.*

**Table S7**. Output for GLMMs predicting within-lake beta-diversity in each system.

|  | *Habitat Type Effect Size* | | *Std. Error* | *z-value* | *Pr(>\|z\|)* | *R_m_^2^* |
| --- | --- | --- | --- | --- | --- | --- |
| ***Sierra Nevada*** | |  |  |  |  |  |
| Within-Lake Temporal Beta | | 0.71 | 0.09 | 7.5 | **4.53x10^-14^** | 0.71 |
| ***Vancouver Island*** | |  |  |  |  |  |
| Within-Lake Spatial Beta | | 2.84 | 0.11 | 24.2 | **< 2 x10^-16^** | 0.99 |

Generalized Linear Mixed Effects Models (GLMMs) within-lake temporal (Sierra Nevada) and within-lake spatial (Vancouver Island) beta diversity. Models were fit with a beta distribution using a logit link and habitat type as a fixed effect. In the Sierra Nevada, lake was a random intercept. For Vancouver Island, lake and year were random intercepts. Columns indicate the effect size of habitat type (logit space), the standard error of the effect size estimate, z-value, P-value, and the marginal R^2^ of models which represents the amount of variation explained by habitat type.

**Table S8.** PERMANOVA balance subsampling sensitivity analysis.

|  | *Dist. Metric* | *Model* | *Full R^2^* | *Median Sens. R^2^* | | *Prop. Sign.* |
| --- | --- | --- | --- | --- | --- | --- |
| ***Sierra Nevada*** |  |  |  |  |  | |
| Habitat Type | Bray-Curtis | Spatial | 0.21 | 0.28 | **1.00** | |
| Lake | Bray-Curtis | Spatial | 0.16 | 0.23 | **1.00** | |
| Habitat Type * Lake | Bray-Curtis | Spatial | 0.10 | 0.22 | **1.00** | |
| Habitat Type | Bray-Curtis | Temporal | 0.21 | 0.28 | **1.00** | |
| Year | Bray-Curtis | Temporal | 0.04 | 0.02 | 0.03 | |
| Habitat Type * Year | Bray-Curtis | Temporal | 0.007 | 0.02 | 0.04 | |
| Habitat Type | weight-Uni | Spatial | 0.39 | 0.49 | **1.00** | |
| Lake | weight-Uni | Spatial | 0.10 | 0.14 | 0.17 | |
| Habitat Type * Lake | weight-Uni | Spatial | 0.07 | 0.14 | 0.14 | |
| Habitat Type | weight-Uni | Temporal | 0.39 | 0.49 | **1.00** | |
| Year | weight-Uni | Temporal | 0.03 | 0.02 | 0.31 | |
| Habitat Type * Year | weight-Uni | Temporal | 0.009 | 0.02 | 0.12 | |
| ***Vancouver Island*** |  |  |  |  |  | |
| Habitat Type | Bray-Curtis | Spatial | 0.09 | 0.22 | **1.00** | |
| Lake | Bray-Curtis | Spatial | 0.11 | 0.20 | **1.00** | |
| Habitat Type * Lake | Bray-Curtis | Spatial | 0.05 | 0.19 | **1.00** | |
| Habitat Type | Bray-Curtis | Temporal | 0.09 | 0.22 | **1.00** | |
| Year | Bray-Curtis | Temporal | 0.03 | 0.04 | **1.00** | |
| Habitat Type * Year | Bray-Curtis | Temporal | 0.01 | 0.04 | **1.00** | |
| Habitat Type | weight-Uni | Spatial | 0.11 | 0.26 | **1.00** | |
| Lake | weight-Uni | Spatial | 0.11 | 0.19 | **1.00** | |
| Habitat Type * Lake | weight-Uni | Spatial | 0.05 | 0.18 | **1.00** | |
| Habitat Type | weight-Uni | Temporal | 0.11 | 0.26 | **1.00** | |
| Year | weight-Uni | Temporal | 0.03 | 0.05 | **1.00** | |
| Habitat Type * Year | weight-Uni | Temporal | 0.01 | 0.04 | **1.00** | |

Balanced subsampling sensitivity analysis to account for unequal sampling sizes in PERMANOVAs between fish and water microbiomes. Full R^2^ values reflect effect sizes of terms when using the full data set. Median sens. R^2^ values are the median effect sizes for terms across 100 iterations of model fitting with equal sample sizes between fish and water samples. Prop. sign. represents the proportion of model iterations were p-values for terms were significant (p < 0.05), terms where 100% of iterations were significant are in bold.

**Table S9.** Balanced subsampling sensitivity analysis for within-lake spatial and temporal beta-diversity.

|  | *Full Habitat Type Est.* | | *Median Perm. Est.* | | *95% Perm. Range* | *Median R_m_^2^* | *Prop. Sign.* |
| --- | --- | --- | --- | --- | --- | --- | --- |
| ***Sierra Nevada*** | |  |  |  | |  |  |
| Temporal Beta | | 0.71 | 0.78 | 0.42 – 1.16 | | 0.69 | 0.99 |
| ***Vancouver Island*** | |  |  |  | |  |  |
| Spatial Beta | | 1.73 | 2.01 | 1.92 – 2.09 | | 0.96 | 1.00 |

Balanced subsampling sensitivity analysis to evaluate whether unequal sample sizes between fish and water influence differences in within-lake beta-diversity between habitat types. The full habitat type estimate column reflects the effect size of habitat type on temporal (Sierra Nevada) and spatial (Vancouver Island) within-lake beta-diversity. Median permutation estimates, the 95% permutation range of estimates, median marginal R^2^ values, and the proportion of significant effects were calculated across the 999 permutations performed.

**Table S10.** Partial Mantel tests examining the independent effects of elevation and geographic distance on Sierra Nevada microbial communities.

| ***Effect of:*** | Geographic Distance | | Elevation Distance | |
| --- | --- | --- | --- | --- |
| ***Controlling for:*** | Elevation Distance | | Geographic Distance | |
|  | *ρ* | *P-value* | *ρ* | *P-value* |
| Brook Trout | -0.11 | **0.001** | -0.15 | **0.001** |
| Sierra Nevada Lakes | 0.06 | **0.02** | -0.21 | **0.001** |

Partial Mantel tests generated from 999 permutations; *ρ = Spearman’s partial mantel correlation. Significant effects (p < 0.05) are in bold.*

**Table S11.** PERMANOVA output table based on Unifrac distances between model partitions for each sample category (brook trout, stickleback, Sierra Nevada bacterioplankton, Vancouver Island bacterioplankton). Permutations were restricted by system (strata = study system).

|  | *df* | *SS* | *R^2^* | *F* | *Pr(>F)* |
| --- | --- | --- | --- | --- | --- |
| Model Partition | 2 | 1.35 | 0.31 | 2.58 | **0.001** |
| Habitat Type | 1 | 0.61 | 0.14 | 2.33 | **0.007** |
| Model Partition*Habitat Type | 2 | 0.89 | 0.20 | 1.71 | **0.028** |
| Residual | 6 | 1.56 | 0.35 |  |  |
| Total | 11 | 4.42 | 1.00 |  |  |

PERMANOVA tables generated from 999 permutations; *df = degrees of freedom; SS = sum of squares. Significant effects (p < 0.05) are in bold.*

**FIGURES**

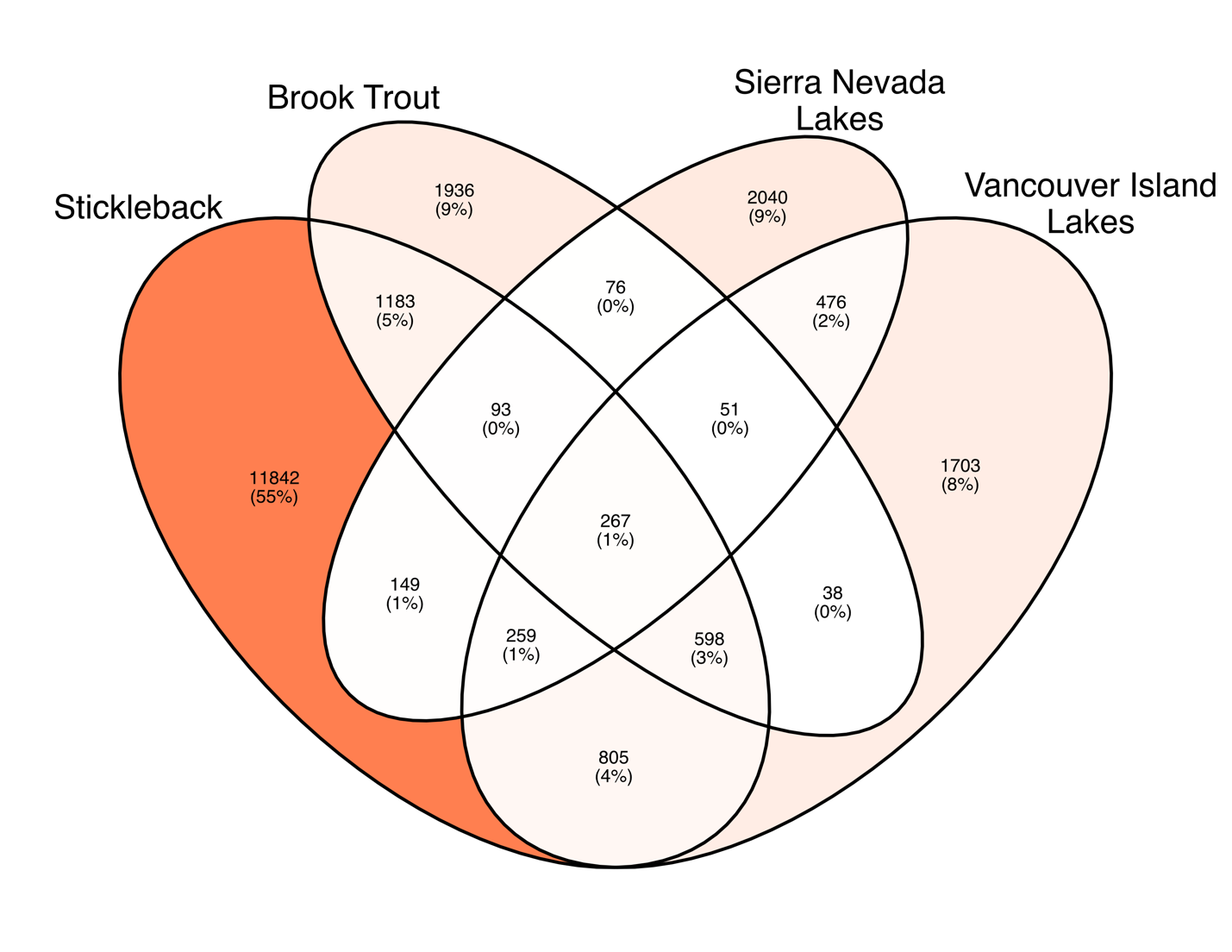

**Fig S1.** Venn diagram of shared ASVs across sample categories.

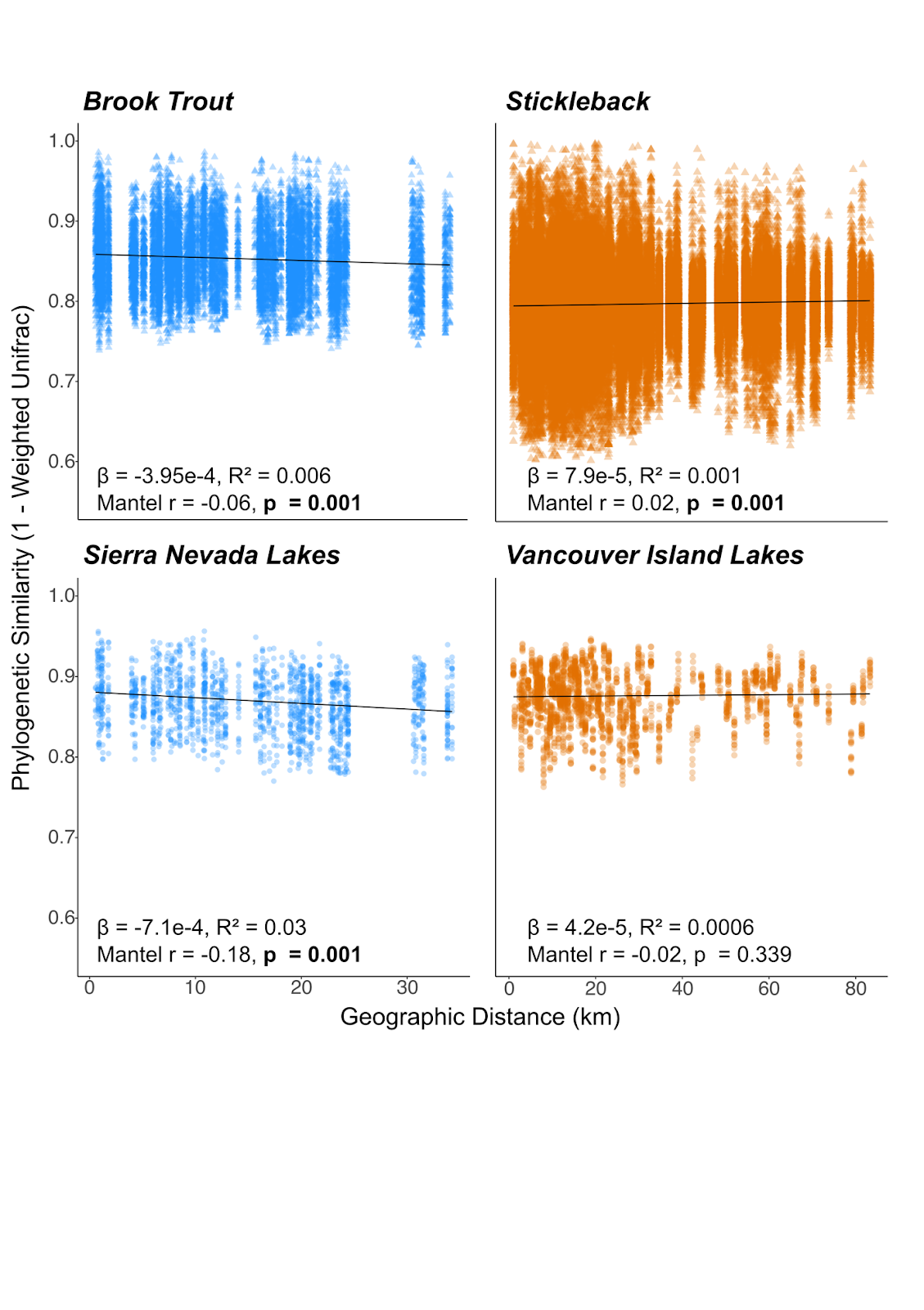

**Figure S2.** Phylogenetic distance decay for each sample category that depicts pairwise phylogenetic microbiome similarity (1 – Weighted-Unifrac distance) as a function of geographic distance. Points are colored by study system (blue – Sierra Nevada, orange – Vancouver Island) and shape denotes sample type (triangle – fish gut, circle – lake water). Slopes and adjusted R^2^ values were calculated from linear mixed effects models with comparisons for communities from the same year as a random effect. Mantel r values and the corresponding p-values are reported for determining the significance of relationships.

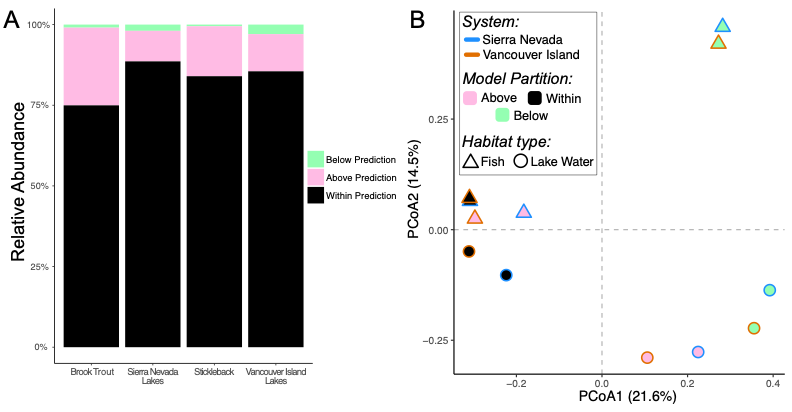

**Figure S3.**  Neutral model outputs. **(A)** Barplot of the proportion of ASVs Above (pink), Below (green), and Within (black) the prokaryote neutral model prediction for each sample category. **(B)** Principle coordinates analysis (PCoA) based on phylogenetic dissimilarity (unifrac distance) between ASVs above, within, and below the confidence interval of the predicted frequency of occurrence for each sample category. Shape denotes the sample type (*triangle* – fish gut, *circle* – lake water) and the fill color corresponds to the model partition (*black* – within, *pink* – above, *green* – below).

rRNA 806R gene primer greatly increases detection of SAR11 bacterioplankton. Aquatic Microbial Ecology, 75(2), 129–137. http://doi.org/10.3354/ame01753

Caporaso JG, Lauber CL, Walters WA, Berg-Lyons D, Huntley J, Fierer N, et al. Ultra-high-

throughput microbial community analysis on the Illumina HiSeq and MiSeq platforms.ISME J 2012; 6: 1621–1624.

Parada, A. E., Needham, D. M., & Fuhrman, J. A. (2016). Every base matters: assessing small

subunit rRNA primers for marine microbiomes with mock communities, time series and global field samples. Environmental Microbiology, 18(5), 1403–1414

Walters, W., Hyde, E. R., Berg-Lyons, D., Ackermann, G., Humphrey, G., Parada, A., Gilbert, J.

A., Jansson, J. K., Caporaso, J. G., Fuhrman, J. A., Apprill, A., & Knight, R. (2016). Improved bacterial 16S rRNA gene (V4 and V4-5) and fungal internal transcribed spacer marker gene primers for microbial community surveys. *mSystems*, *1*(1). https://doi.org/10.1128/mSystems.00009-15
